# Disbalance of the intestinal epithelial cell turnover and apoptosis in a rat model of sporadic Alzheimer’s disease

**DOI:** 10.1101/2021.04.22.440947

**Authors:** Jan Homolak, Ana Babic Perhoc, Ana Knezovic, Jelena Osmanovic Barilar, Fatma Koc, Catherine Stanton, R. Paul Ross, Melita Salkovic-Petrisic

## Abstract

**Background:** Dyshomeostasis of the gastrointestinal (GI) system is investigated as a potential contributor to metabolic dysfunction, systemic and neuro-inflammation recognized as important pathophysiological drivers of neurodegeneration. Gastrointestinal redox dyshomeostasis and dysfunctional brain-gut incretin axis have been reported in the rat model of insulin-resistant brain state (IRBS)-driven neurodegeneration induced by intracerebroventricular administration of streptozotocin (STZ-icv). The aim was to assess i) whether GI oxidative stress is accompanied by structural and functional changes of the epithelial barrier; ii) whether the brain glucose-dependent insulinotropic polypeptide receptor (GIP-R) is also involved in redox regulation of the gut; and iii) whether the STZ-icv brain-gut axis is resistant to pharmacological inhibition of the brain GIP-R.

**Methods:** Forty three-month-old male Wistar rats were treated with 3mg/kg STZ-icv or vehicle. One month later the animals were randomized to receive either saline or 85 μg/kg GIP-R inhibitor [Pro^3^]-GIP intracerebroventricularly and sacrificed 30 minutes later. Thiobarbituric acid reactive substances (TBARS) were measured in plasma and duodenum. Duodenal sections were subjected to morphometric analysis. Caspase-3 expression and activation were analyzed by western blot and spatial signal analysis was done by multiplex fluorescent signal amplification (MFSA). Data were analyzed by linear and linear mixed modeling, and exploration was done by principal component analysis.

**Results:** Inhibition of the brain GIP-R decreased plasma TBARS in the controls and the STZ-icv animals and increased duodenal TBARS only in the controls. Acute inhibition of brain GIP-R affects duodenal epithelial cell, but not villus structure, while all morphometric parameters were altered in the STZ-icv-treated animals. Morphometric changes in the STZ-icv animals were accompanied by reduced levels of activated and total regulator of apoptosis – caspase-3. Acute inhibition of brain GIP-R inactivated duodenal apoptosis at the level of caspase-3 activation.

**Conclusions:** Brain GIP-R is involved in the regulation of the systemic and duodenal redox homeostasis and epithelial function. Duodenal oxidative stress in the STZ-icv rats is accompanied by the resistance of the brain-gut GIP axis and morphological changes indicative of abnormal epithelial cell turnover and dysfunctional GI barrier. Dysfunction of the brain-gut incretin axis might be an important etiopathogenetic factor in neurodegeneration and a potential pharmacological target.

## Introduction

Alzheimer’s disease (AD) is the most common neurodegenerative disorder characterized by progressive cognitive impairment. The prevalence rates of AD rise exponentially with age and predictions suggest that its global burden will continue to rise with the increasing median age of the population[1]. The mendelian inheritance of amyloid precursor protein, presenilin-1 or presenilin-2 seems to be the main driver of the disease in a small fraction of patients (~5%), however, the etiopathogenesis of the sporadic form of AD (sAD) accounting for the residual ~95% is still unclear [2].

The gastrointestinal (GI) system is increasingly recognized as an important player involved in the pathophysiology of sAD. A recent systematic review and meta-analysis suggested GI disorders are increasingly prevalent in AD and Parkinson’s disease patients [3], and accumulating evidence suggests brain neurodegenerative processes are accompanied by alterations of gut microbiota[4,5]. Although the gut and brain dyshomeostasis in neurodegeneration seem to be closely related, it is currently unknown whether the prodromal GI symptoms are a consequence of early changes in the brain or emerge as an independent pathophysiological entity. The brain regulates permeability, motility, secretion, and the immune function of the gut via the efferent arm of the autonomic nervous system-mediated modulation of the enteric nervous system, smooth muscle contractility, and mucus production and secretion[6]. Furthermore, it has been shown that the central nervous system (CNS) plays a critical role in host-enteric microbiota signaling[7]. Consequently, impaired brain homeostasis in the early presymptomatic phase of AD may trigger pathophysiological processes in the gut. The circadian rhythm dysfunction occurs in the early stage of sAD[8], and host circadian dysrhythmia has been shown to induce dysbiosis and impair intestinal function[9,10]. Furthermore, presymptomatic pathophysiological processes in the CNS may influence GI function and microbiota by affecting the volume, viscosity, and porosity of mucus which plays an integral part in the maintenance of the GI barrier [11]. Interestingly, it has been proposed that brainstem dysfunction precedes neuropathological changes of the supratentorial regions in the early stage of AD [12], and non-invasive measurement of brainstem vagus, the most important neural bidirectional pathway conveying information between the viscera and the brain has been proposed for detection of subtle functional disturbances that may develop even decades before structural damage and clinical symptoms[13]. Once the GI barrier is breached, luminal microbial amyloid and pro-inflammatory molecules induce systemic inflammation and metabolic dysfunction further encouraging the development of insulin-resistant brain state (IRBS) and neurodegeneration [4,14–17]. Furthermore, a dysfunctional GI barrier might enable the seeding of the intestinal amyloid to the CNS via vagus-mediated transport [18]. Taken all together current evidence suggests the dyshomeostasis of the GI system in AD might play an important role in the self-perpetuating vicious cycle of systemic and neuro-inflammation, and may thus be an attractive diagnostic and therapeutic target.

Pathophysiological changes of the GI system have been observed in animal models of AD. The dysregulated gut homeostasis has been reported in transgenic models of AD such as Tg2576, TgCRND8, 5×FAD, mThy1-hAβPP751, AβPP23, APP/PS1 [19–23]. In the Tg2576 model, the observed dyshomeostasis of the GI system occurs early in the presymptomatic phase of the disease, before the accumulation of brain Aβ and the development of cognitive deficits [19]. Moreover, the involvement of the GI tract was characterized by a reduced expression of an important apical tight junction protein E-cadherin[19]. Furthermore, a widespread bacterial breach through the intestinal barrier and the corresponding antigenic load onto the intestinal epithelium and increased concentration of proinflammatory and angiogenic plasma cytokines have been reported [19]. Sohrabi et al. recently demonstrated that gut inflammation induced by administration of dextran sulfate sodium can exacerbate the deposition of Aβ plaques and alter microglial immunoreactivity in the AppNL-G-F mouse model of AD [24]. In contrast to the accumulating evidence related to the intestinal pathology in the transgenic models of AD, the involvement of the GI system in the non-transgenic models of the disease is still unexplored. Apart from our recent report on the duodenal oxidative stress, and resistance of the brain-gut glucagon-like peptide-1 axis in the rats treated intracerebroventricularly with streptozotocin (STZ-icv) [25] we are not aware of any reports on the structure and function of the GI system in the non-transgenic animal models of AD. The non-transgenic models of sAD provide a unique insight into pathomechanisms mediating the effects of risk factors on the progression of the disease and provide an invaluable platform for understanding possible mechanisms implicated in the etiopathogenesis.

The present aim was to further explore the pathophysiological changes in the duodenum of the STZ-icv rats. Considering the GI barrier provides protection against systemic and neuro-inflammation, and retrograde transport of toxic intestinal Aβ to the brain, exploration of the mechanisms that enable functional maintenance of the barrier – namely intestinal epithelial cell turnover and apoptosis were the main focus of our research. Streptozotocin is a nitrosourea compound used for modeling insulin resistance and diabetes mellitus type 1 in rodents following parenteral administration [26,27]. After intracerebroventricular administration, streptozotocin has been shown to cause cerebral pathophysiological changes resembling those found in sAD such as the insulin-resistant brain state (IRBS) [28,29], glucose hypometabolism [29,30], oxidative stress [31], mitochondrial dysfunction [28], neuroinflammation [32,33], cholinergic deficits [34], accumulation of amyloid β (Aβ) [35] and hyperphosphorylated tau [36], and the development of cognitive deficits [37].

Recently, a dysfunctional incretin system was also reported in STZ-icv rats. An active fraction of plasma glucagon-like peptide-1 (GLP-1) is decreased in the STZ-icv rats 3 months after the treatment (a period corresponding to the early stage of the disease) [30]. Furthermore, GI redox homeostasis was resistant to pharmacological inhibition of the brain GLP-1 receptors (GLP-1R) in the STZ-icv rats indicating incretin resistance of the brain-gut axis 1 month after the model induction [25]. Incretins are emerging as important players in neurodegeneration both as factors involved in the etiopathogenesis, and attractive pleiotropic therapeutic targets as they can modulate IRBS, oxidative stress, Aβ homeostasis, apoptosis, cell proliferation and differentiation, synaptic function, and peripheral metabolism [38–43]. Recent evidence suggests that incretin hormone glucose-dependent insulinotropic polypeptide (GIP) exerts strong neuroprotective properties and that it might be suitable for targeting IRBS in neurodegeneration alone or in combination with GLP-1[44,45]. Consequently, we wanted to explore whether brain GIP receptors (GIP-R) are involved in the regulation of intestinal redox homeostasis and whether the STZ-icv rats develop resistance of the brain-gut GIP axis similarly as has been shown for GLP-1[25].

## Materials and methods

### Animals

Three-month-old male Wistar rats (n=40) from the Department of Pharmacology (University of Zagreb School of Medicine) were used in the experiment. The animals were kept 2-3/cage with standardized pellets and water available *ad libitum*. The light-dark cycle was 7AM/7PM, the humidity was 40-70%, and the temperature 21-23°C. Standard bedding (Mucedola S.R.L., Italy) was changed twice per week.

### Streptozotocin treatment

STZ-icv was used to model IRBS as described previously [30,33]. Rats were randomly assigned to either the control group (CTR; n=20) or the streptozotocin group (STZ; n=20). Each animal was anesthetized with ketamine (70 mg/kg) and xylazine (7mg/kg), the skin was opened and the skull was trepanated bilaterally with a high-speed drill. The treatment consisted of bilateral intracerebroventricular administration (2μl/ventricle) repeated two times with an inter-treatment interval of 48 hours. The control group received 0.05M citrate buffer (pH 4.5) on both occasions. The STZ group received streptozotocin dissolved in 0.05M citrate buffer (pH 4.5) in a cumulative dose of 3 mg/kg (1.5 mg/kg followed by 1.5 mg/kg after 48 hours). A modification of the protocol first proposed by Noble et al. was used [46] as described in [47]. The coordinates used were: −1.5 mm posterior; ± 1.5 mm lateral; +4 mm ventral from pia mater relative to bregma.

### Pharmacological inhibition of brain GIP receptors

Brain GIP receptors were acutely inhibited by intracerebroventricular administration of 85 μg/kg GIP receptor antagonist [Pro^3^]-GIP dissolved in saline one month after the STZ-icv administration. The animals from both the CTR and the STZ group were randomized to receive either vehicle (saline) or [Pro^3^]-GIP by a single bilateral intracerebroventricular injection (2μl/ventricle).

### Sacrifice and tissue collection

Thirty minutes after intracerebroventricular treatment with either [Pro^3^]-GIP (GIPI) or saline, 6/10 animals from each group were euthanized in general anesthesia and decapitated. The post-gastric 2 cm of proximal duodenum was dissected, cleared from the surrounding tissue, and luminal content in ice-cold phosphate-buffered saline (PBS). The tissue was snap-frozen in liquid nitrogen and stored at −80 °C. The samples were homogenized on dry ice and subjected to three cycles of sonification (Microson Ultrasonic Cell 167 Disruptor XL, Misonix, SAD) in five volumes of lysis buffer containing 150 mM NaCl, 50 mM Tris-HCl pH 7.4, 1 mM EDTA, 1% Triton X-100, 1% sodium deoxycholate, 0.1% SDS, 1 mM PMSF, protease inhibitor cocktail (Sigma-Aldrich, USA) and phosphatase inhibitor (PhosSTOP, Roche, Switzerland) (pH 7.5) on ice. Homogenates were centrifuged for 10 minutes at a relative centrifugal force (RCF) of 12879 × g and 4°C. The supernatant protein concentration was assessed by quantification of the change in absorbance of the Bradford reagent (Sigma-Aldrich, USA) at 595 nm utilizing Infinite F200 PRO multimodal microplate reader (Tecan, Switzerland). Samples were stored at −80 °C until further analysis. Plasma was extracted from whole blood drawn from the retro-orbital sinus after centrifugation at 1159 × g RCF at 4°C for 10 minutes in heparinized tubes (100 μl/sample). Four animals from each group were subjected to transcardial perfusion with 4% paraformaldehyde (pH 7.4), the tissue was removed, equilibrated in paraformaldehyde and after equilibration, each intestinal segment was cut into approximately 8 equal subsegments and embedded in a paraffin block to provide 8 transverse sections corresponding to different layers in each section.

### Lipid peroxidation

Thiobarbituric acid reactive substances (TBARS) assay was used for assessment of lipid peroxidation as described previously [25,48,49]. Briefly, tissue homogenate supernatant or plasma (12 μl) were mixed with 120 μl TBA-TCA reagent (0,375% thiobarbituric acid in 15% trichloroacetic acid) and 70 μl of ddH_2_O. Samples were placed in perforated microcentrifuge tubes and incubated for 20 minutes in a heating block set at 95°C. The thiobarbituric acid-malondialdehyde complex was extracted in 220 μl n-butanol. The absorbance of the butanol fraction was analyzed at 540 nm in a 384-well plate using an Infinite F200 PRO multimodal microplate reader (Tecan, Switzerland). Malondialdehyde (MDA) concentration was assessed with a linear model using a dilution curve of MDA tetrabutylammonium salt in ddH_2_O.

### Morphometric analyses

The morphometric analysis was done on tissue sections treated with 4′,6-diamidino-2-phenylindole (DAPI). The paraffin blocks were cut on a microtome to obtain 5 μm thin sections representing ~8 subsegments each. Slides were then deparaffinized by a standard protocol through xylene and a sequence of solutions with a decreasing concentration of ethanol (EtOH). Slides were equilibrated in PBS and coverslipped with the Fluoroshield mounting medium containing DAPI. A U-MNU2 filter set (EX: 365/10; EM: >420), epifluorescence microscope Olympus BX51, and CellSens Dimensions software were used to obtain epifluorescent images of tissue sections that were subsequently analyzed in Fiji (NIH, USA). Morphometric analyses were done on a dataset of random sample fluorescent images to reduce bias. If obvious artifacts were present, the closest neighboring structure was sampled. Villus-crypt pairs were measured using a segmented line technique. Villus length and crypt depth ratios (VL/CD) were calculated for each villus-crypt pair and a sample of at least 8 random pairs were obtained for each animal. Epithelial cell height (ECH) and epithelial cell width (ECW) were calculated from a random sample of villus tip epithelial cells. For each cell sampled, ECH was measured as the distance from the center of the nucleus to the luminal border of the cell, and ECW was calculated as the distance from the center of the nucleus to the center of the nucleus of the closest neighboring enterocyte.

### Western blot

Western blot was used to analyze caspase-3 in tissue homogenates. Briefly, tissue samples were mixed with a standard sample buffer in a 2:1 volumetric ratio, heated at 95 °C, spun down, and the volumes corresponding to 35 μg were loaded onto TGX Stain-Free FastCast 12% polyacrylamide gels (Bio-Rad, USA) for electrophoretic separation. The separated proteins were visualized using ChemiDoc MP Imaging System UV transilluminator (Bio-Rad, USA) for subsequent total protein correction. The proteins were transferred onto the nitrocellulose membranes using Trans-Blot Turbo semi-dry transfer protocol (2.5A; 25V; 7 min). Protein fixation and residual peroxidase activity inhibition were done by incubation in 1% Ponceau S in 5% acetic acid[50]. Membranes were washed for 5 min in LSWB, blocked for 1 hour at room temperature (RT) in a blocking buffer (5% nonfat dry milk solution with 0,5% Tween 20 in low-salt washing buffer (LSWB: 10 mM Tris, 150 mM NaCl, pH 7.5)), and incubated with 1:500 anti-caspase-3 antibody (#9662S, Cell Signaling Technologies, USA) overnight at 4 °C in-between polyvinyl chloride (PVC) sheets to ensure equal reagent dispersion. Membranes were washed 3×5 min in LSWB, incubated with 1:2000 anti-rabbit secondary antibody (#7074S, Cell Signaling Technologies, USA) for 1 h at RT, washed 3×5 min in LSWB again, and visualized using chemiluminescent reagent (SuperSignal™ West Femto; Thermo Fisher Scientific, USA). Luminescence was captured by MicroChemi high-performance chemiluminescence imager (DNR Bio-Imaging Systems Ltd., Israel) and GelCapture software. Immunoblot densitometry and total protein quantification were done in Fiji (NIH, USA) using Gel Analyzer protocol after Rolling Ball background subtraction. Raw western blot data provided in **Supplement 1**.

### Antigen retrieval and elimination of autofluorescence in formalin-fixed paraffin-embedded tissue sections

Paraffin blocks were cut on a microtome to obtain 5 μm thin sections representing ~8 subsegments each. Slides were then deparaffinized by a standard protocol through xylene and a sequence of solutions with a decreasing concentration of EtOH. The tissue sections were secured in between two histological slides using custom-made spacers and insulating tape (not covering tissue sections) and placed in an antigen retrieval solution (10mM sodium citrate, 0.05% Tween 20, pH 6.0) at 95 °C for 1 h. Tissue sections were removed from the retrieval solution, briefly washed in ddH_2_O, and placed in a container filled with 0.05% sodium azide in PBS in a custom-made autofluorescence-quenching apparatus to reduce fixative-induced and cellular-derived artifactual autofluorescence. A 300 W full spectrum light-emitting diode hydroponic panel was used as a constant light source and tissue sections were illuminated in azide-PBS for 48 hours[51,52].

### Multiplex fluorescent signal amplification

Ultrasensitive detection of cytoplasmic and nuclear tissue caspase-3 was conducted utilizing immunohistochemistry followed by a combination of sensitive avidin-biotin-based amplification and catalyzed reporter deposition. Biotinylated secondary antibodies were used for avidin-biotin-complex-based specific deposition of biotinylated horseradish peroxidases (bHRP) in the epitope proximity. Catalyzed reporter deposition was then employed to exploit epitope-bound bHRP complexes for the generation of tyramide radicals and covalent deposition of fluorescent tyramide conjugates to proximal tyrosine residues.

### Immunohistochemistry and avidin-biotin amplification

Following autofluorescence quenching, insulating tape and protective glass slides were removed, tissue was briefly washed in ddH_2_O and equilibrated in PBS. In all subsequent steps, PVC coverslips were used to ensure the optimal distribution of the applied reagents on the slides. Residual peroxidases were blocked by incubation in 1% H_2_O_2_ in PBS for 15 min. Sections were washed for 5 min in PBS and incubated in a blocking solution for 1 h (5% normal goat serum (NGS) in 0.25% Triton X-100 in PBS (PBST)). Sections were incubated in anti-caspase-3 primary antibody (#9662S, Cell Signaling Technologies, USA) diluted 1:500 in 1% NGS in PBST overnight at 4 °C under PVC coverslips. Slides were washed 3×5 min in PBS and incubated with 1:1000 anti-rabbit biotinylated secondary antibody (BA-1000, Vector Laboratories, USA) in 1% NGS in PBST for 1 h at RT. Slides were washed 3×5 min in PBS and incubated in freshly prepared avidin-bHRP solution (Vector Laboratories, USA) for 30 min at RT. Sections were washed 3×5 min in PBS and subjected to catalyzed reporter deposition amplification.

### Catalyzed reporter deposition (CARD) amplification

Localization of caspase-3 in the tissue was done by catalyzed reporter deposition amplification utilizing fluorescent tyramide conjugates. Reporter synthesis was done as follows: 1 mg of tyramine-HCl was dissolved in 100 μl of dimethylformamide (DMF) and 1 μl of 7.2 M triethylamine (TEA) (1.25-fold equimolar) was added for tyramine deprotonation. 34 μl of the solution were mixed with 100 μl of the freshly dissolved amine-reactive N-succinimidyl ester of water-soluble stable fluorophore Atto488 in dimethylformamide (1 mg/100 μl). The solution was mixed thoroughly and left to react for 2 h at RT. After 2 h, 867 μl of EtOH was added to the solution to obtain the 1 mg/ml haptenized tyramide stock[53].

The haptenized tyramide stock was mixed with a freshly prepared reaction buffer (0.001% H_2_O_2_ in 0.1M imidazole in PBS) in a 1:100 ratio before the 3×5 min PBS washing step following the incubation with the avidin-bHRP solution to make the tyramide reaction solution (TRS). After the 3×5 min PBS washing step, sections were incubated with the TRS for 15 min and washed for 5 min in PBS. Sections were incubated in the Sudan Black B solution (0.3% in 70% EtOH) for 10 minutes for an additional improvement of the signal-to-noise ratio and the leftover dye was destained with 70% EtOH. Tissue sections were equilibrated in PBS and coverslipped with Fluoroshield aqueous mounting medium containing DAPI.

### Image analysis

Image analysis was done in Fiji (NIH, USA). Epifluorescent images were obtained utilizing Olympus BX51 and CellSens Dimensions software. U-MNU2 and U-MNIB2 filter sets were used for signal acquisition and U-MWIG2 was used as an internal control for residual autofluorescence. U-MNU2-obtained images corresponding to DAPI were used to calculate nuclear masks used for further segmentation. Masks were generated by pixel-based segmentation utilizing trainable weka segmentation machine learning algorithms[54]. Fill Holes and Watershed algorithms were used for post-processing and the masks were redirected to the U-MNIB2 dataset pre-processed by rolling ball background subtraction. 100 pixels were used as the initial particle analysis cut-off, and further filtering was done in R. The nuclear signal >650 pixels was used for subsequent analyses as it was found to be an optimal cut-off point by iterative nuclear segmentation. Signal intensity analysis was done based on integrated density measurements. A red lookup table was used for the visualization of the U-MNU2 signal.

### Data analysis

Data were analyzed in R (4.0.2). Overall effects of treatment 1 (TH1: intracerebroventricular citrate buffer vs. STZ) and treatment 2 (TH2: intracerebroventricular saline vs. [Pro^3^]-GIP) were analyzed by linear regression using TH1 and TH2 as independent variables. Moderation of the STZ-icv effect was assessed by including the treatment interaction term. For oxidative stress assessment, TBARS was used as the dependent variable. For western blot analysis of caspase-3, the total protein-corrected caspase-3 chemiluminescent signal was used as the dependent variable. Linear mixed models were used for the analysis of morphometric data and caspase-3 immunofluorescence to account for the hierarchical structure of the data. The group-pooled raw data pertaining to individual cells was presented alongside the models. TH1 and TH2 were modeled as fixed effects and animal ID as a random effect. Subsequent moderation analysis was done by including the TH1:TH2 interaction in the model. Differences of estimated marginal means or ratios (for log-transformed dependent variables) and respective 95% confidence intervals were reported. P-values were reported only for the main effects models and the interaction terms. The association between the intestinal morphometric parameters and activation of the epithelial caspase-3 was analyzed by principal component analysis. The contribution of individual variables and the individual animals were reported for the calculated biplot. Additional information regarding the principal component analysis is available in **Supplement 2**.

## Results

### The effects of the acute pharmacological inhibition of the brain GIP-R on systemic and duodenal oxidative stress

Acute pharmacological inhibition of endogenous GIP-R signaling utilizing intracerebroventricular administration of [Pro^3^]-GIP decreased the concentrations of lipid peroxidation marker TBARS in plasma of the control and STZ-icv rats (**Fig 1A-C**). In contrast, the same treatment induced duodenal oxidative stress reflected by an increased concentration of lipid peroxidation end products in the control animals (**Fig 1D-F**). Interestingly, the effect was absent in the rat model of sAD indicating a possible resistance of the brain-gut incretin-mediated redox regulation (**Fig 1D-F**).

**Fig 1.**
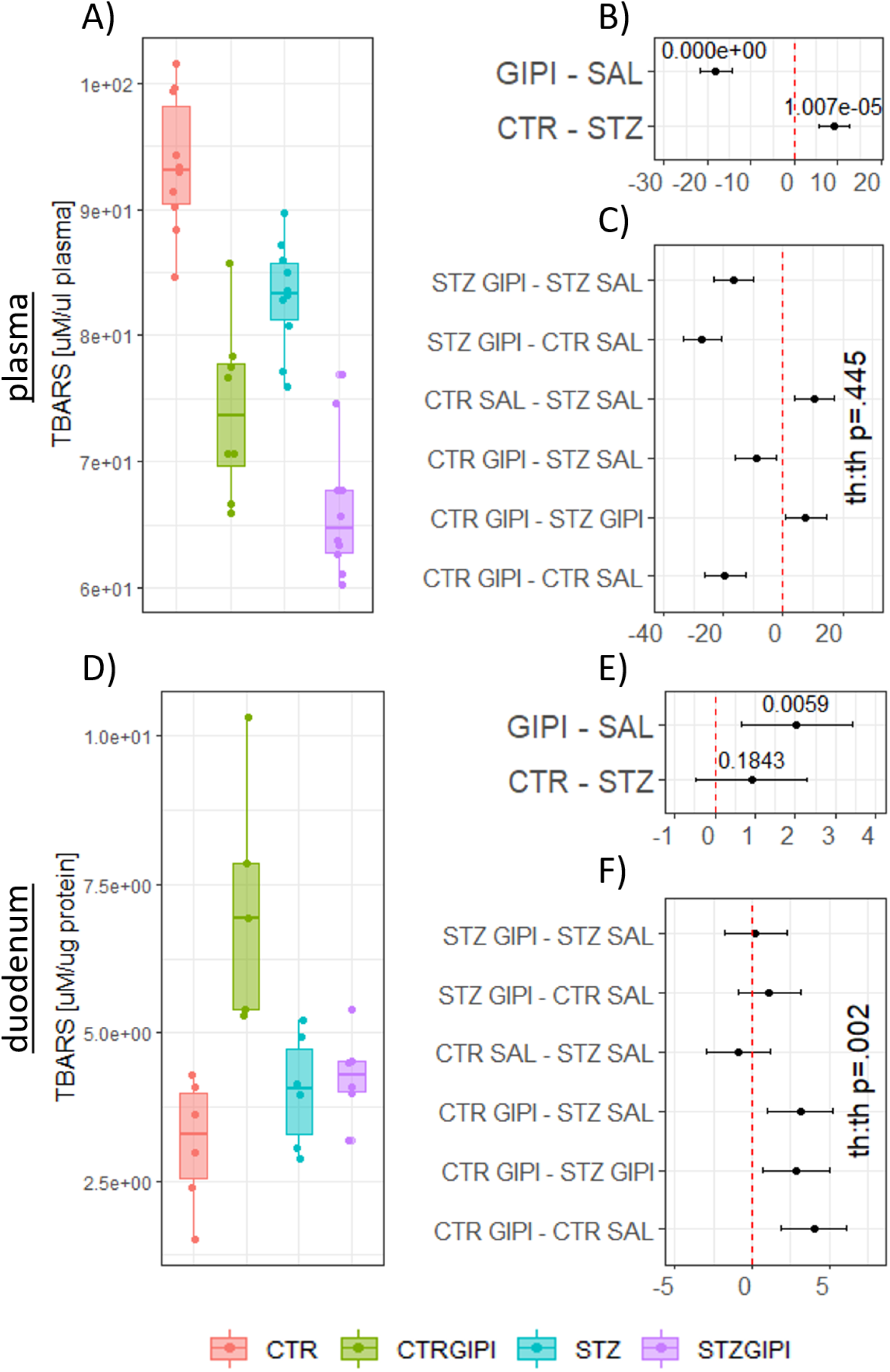
The effect of intracerebroventricular glucose-dependent insulinotropic polypeptide receptor inhibitor [Pro^3^]-GIP (GIPI) on oxidative stress in plasma and duodenum in control rats and the rat model of sporadic Alzheimer’s disease induced by intracerebroventricular streptozotocin (STZ). **(A)** Raw data of plasma thiobarbituric reactive substances (TBARS) reflecting systemic oxidative stress. The effects of STZ and GIPI on plasma TBARS are reported as differences of estimated marginal means extracted from **(B)** the main effects model or **(C)** taking into account treatment interaction. **(D)** Raw data of duodenal thiobarbituric reactive substances (TBARS) reflecting tissue stress. The effects of STZ and GIPI on duodenal TBARS are reported as differences of estimated marginal means extracted from **(E)** the main effects model or **(F)** taking into account treatment interaction. CTR – control animals; STZ – animals treated intracerebroventricularly with streptozotocin; CTRGIPI – control animals treated intracerebroventricularly with GIPI; STZGIPI – STZ animals treated intracerebroventricularly with GIPI.

### Morphometric analysis reveals abnormal duodenal epithelial cell turnover in the STZ-icv rats

In the morphometric analysis, we focused on villus length (VL), crypt depth (CD), VL and CD ratio (VL/CD), epithelial cell height (ECH), and width (ECW). Previous research has shown that VL reflects intestinal homeostasis and function, and VL/CD provides an insight into epithelial cell turnover that is indispensable for the maintenance of intestinal barrier and nutrient absorption [55]. Epithelial cell morphology reflects active mechanisms involved in reepitelization following epithelial shedding as flattening of the remaining cells ensures the protection of the denuded basement membrane during restitution [56]. All three villus-related morphometric parameters were affected in the STZ-icv animals with changes indicative of epithelial turnover disbalance (**Fig 2A-C**). VL/CD and VL were reduced, and CD was increased in the STZ-icv animals (**Fig 2A-C**). Villus tip epithelial cells were shorter in the STZ-icv rats in comparison with the controls, and their height-to-width ratio was reduced (**Fig 2D,E**). The horizontal distance between adjacent cell nuclei was increased in the rat model of sAD (**Fig 2F**).

**Fig 2.**
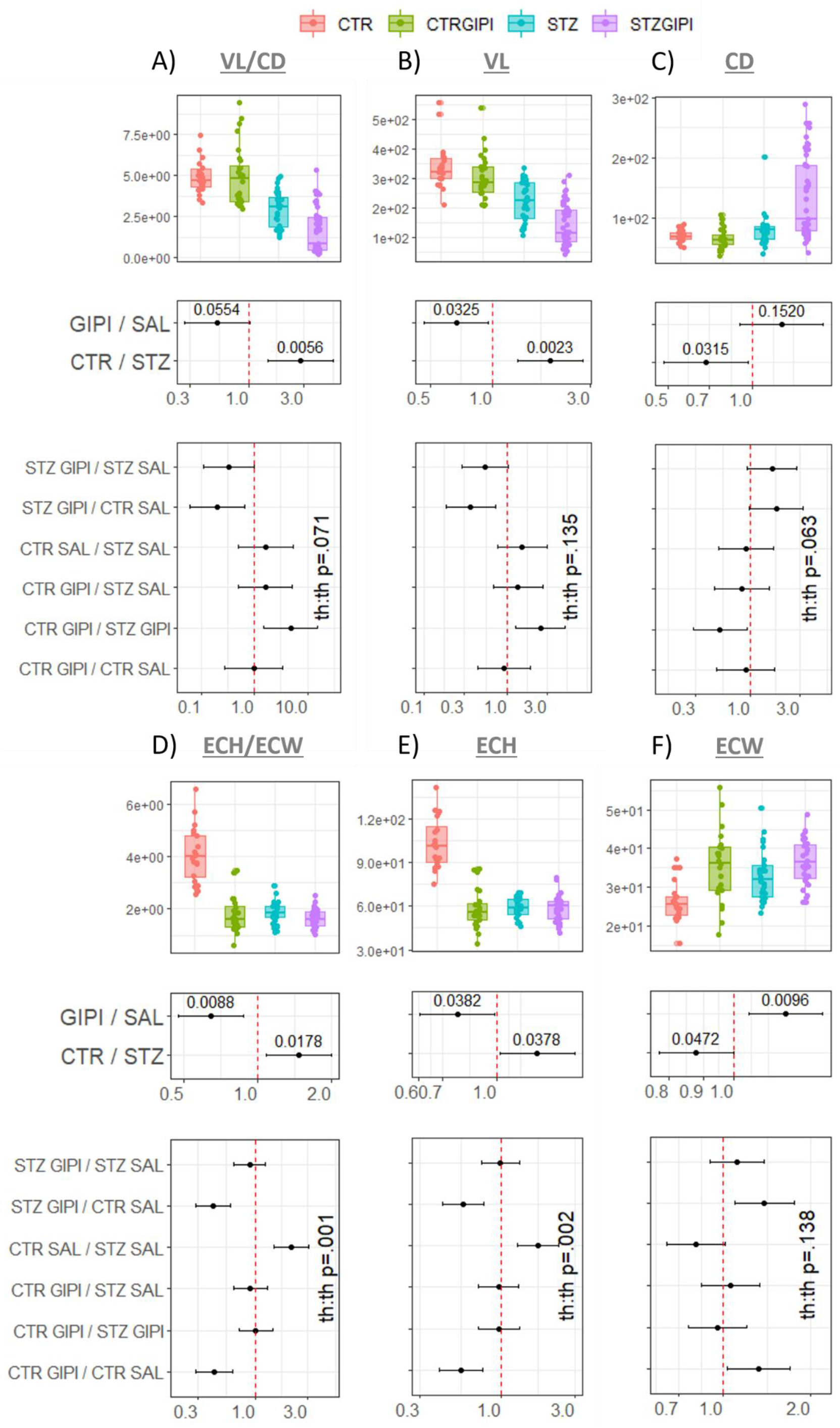
Villus and villus tip epithelial cell morphometric parameters in control rats and the rat model of sporadic Alzheimer’s disease induced by intracerebroventricular streptozotocin (STZ-icv) with and without acute pharmacological inhibition of the brain glucose-dependent insulinotropic polypeptide receptor with [Pro3]-GIP. **(A)** Raw data, main effects, and the interaction mixed-effect model with villus length to crypt depth ratio (VL/CD) used as the dependent variable and treatments as independent variables. **(B)** Raw data, main effects, and the interaction mixed-effect model with villus length (VL) used as the dependent variable and treatments as independent variables. **(C)** Raw data, main effects, and the interaction mixed-effect model with crypt depth (CD) used as the dependent variable and treatments as independent variables. **(D)** Raw data, main effects, and the interaction mixed-effect model with the ratio of villus tip epithelial cell height and width (ECH/ECW) used as the dependent variable and treatments as independent variables. **(E)** Raw data, main effects, and the interaction mixed-effect model with the villus tip epithelial cell height (ECH) used as the dependent variable and treatments as independent variables. **(F)** Raw data, main effects, and the interaction mixed-effect model with the villus tip epithelial cell width (ECW) used as the dependent variable and treatments as independent variables. CTR – control animals; STZ – animals treated intracerebroventricularly with streptozotocin; CTRGIPI – control animals treated intracerebroventricularly with GIPI; STZGIPI – STZ animals treated intracerebroventricularly with GIPI.

### Acute pharmacological inhibition of the brain GIP-R differently affects intestinal morphometric parameters in the controls and the STZ-icv animals

Acute inhibition of the brain GIP-R with intracerebroventricular administration of [Pro^3^]-GIP did not affect villus morphology in the control animals but decreased VL and VL/CD, and increased CD in the rat model of sAD (**Fig 2A-C**). Morphology of the villus tip columnar epithelium was affected by [Pro^3^]-GIP treatment. ECH and ECH/ECW ratio were reduced, and ECW was increased in the controls upon inhibition of the brain GIP-R (**Fig 2D-F**). A similar morphometric pattern was observed in the STZ-icv at baseline, and [Pro^3^]-GIP failed to exert a further reduction of the ECH and ECH/ECW (**Fig 2D,E**).

### The expression and activation of duodenal epithelial caspase-3 is reduced in the rat model of sAD

The expression and activation of the caspase-3, one of the key enzymes involved in the execution phase of cell apoptosis was examined as apoptosis plays an important role in the regulation of intestinal epithelial cell turnover. The western blot analysis of caspase-3 indicated that both the activated nuclear 17 kDa fragment and inactive 30 kDa form were reduced in the duodenal homogenates of the STZ-icv rats (**Fig 2A,B**). Interestingly, the active-to-inactive caspase-3 ratio was largely unchanged between the control and the STZ-icv animals (**Fig 2C**). Multiplex fluorescent signal amplification of the caspase-3 signal in the duodenal epithelium also suggested reduced caspase-3 expression resulting in decreased activation in the STZ-icv animals (**Fig 2D,E**).

### Acute pharmacological inhibition of the brain GIP-R reduces activation of caspase-3 in the duodenal epithelium

Acute pharmacological inhibition of the brain GIP-R decreased the concentration of the activated duodenal caspase-3 in the control and the STZ-icv animals (**Fig 3A**), while the inactive form remained unchanged by the treatment (**Fig 3B**). The active-to-inactive caspase-3 ratio was reduced by inhibition of brain GIP-R in both groups (**Fig 3C**). Multiplex fluorescent signal amplification of the duodenal epithelium indicated no significant change induced by intracerebroventricular [Pro^3^]-GIP in either group (**Fig 3D,E**). The duodenal epithelium was the main focus of our research, however, we also observed a strong caspase-3 signal in a group of subepithelial cells in the control animals upon GIPI treatment (**Fig 3E**– dotted arrow). The exact type of subepithelial cells and the true meaning of this effect remain to be explored. Interestingly, in the STZ-icv animals, most of the remaining caspase-3 signal in the epithelium was localized in cells resembling enteroendocrine cells (**Fig 3E**– arrow).

**Fig 3.**
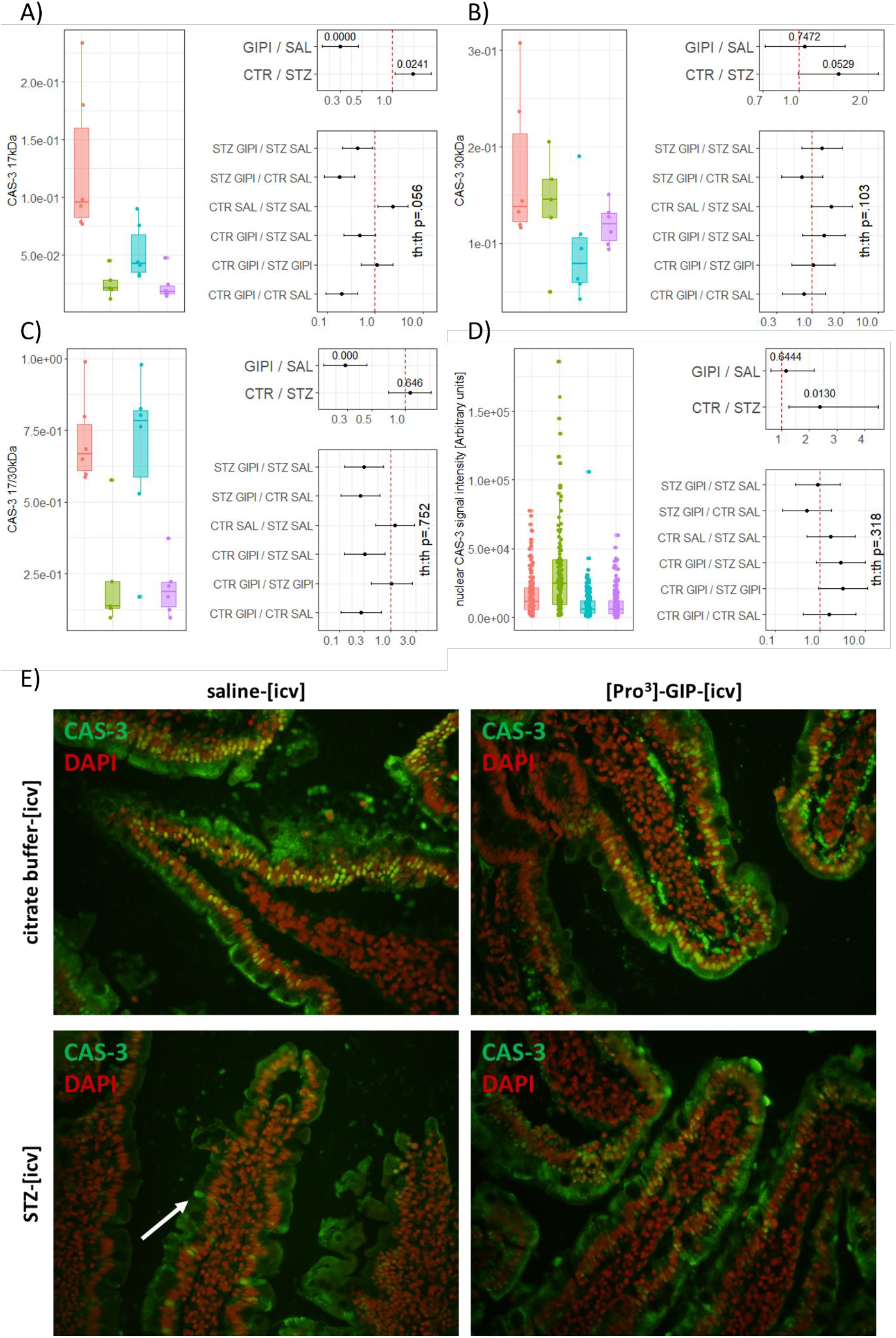
The effect of intracerebroventricular glucose-dependent insulinotropic polypeptide receptor inhibitor [Pro^3^]-GIP (GIPI) on duodenal expression and activation of caspase-3 in control rats and the rat model of sporadic Alzheimer’s disease induced by intracerebroventricular streptozotocin (STZ-icv). **(A)** Western blot analysis of 17 kDa product of caspase-3 corresponding to activated fraction released from an inactive 30 kDa zymogen in the duodenal homogenates. **(B)** Western blot analysis of 30 kDa caspase-3 inactive zymogen in the duodenal homogenates. **(C)** The ratio of activated (17 kDa) to inactive (30 kDa) fraction of caspase-3 in the duodenal homogenates. **(D)** Quantification of the nuclear signal from epifluorescent images. Trainable Weka segmentation machine learning-based segmentation of the 4′,6-diamidino-2-phenylindole nuclear signal was used to generate masks for measurement of the nuclear integrated density of biotinylated horseradish peroxidase-deposited Atto488-haptenized tyramide indicating presence of activated caspase-3 17 kDa fragment. **(E)** Multiplex fluorescent signal amplification of total caspase-3 in the duodenum. The cytoplasmic signal corresponds to the inactive 30 kDa form. The nuclear signal corresponds to the activated 17 kDa fragment of caspase-3. Dotted arrow – a strong caspase-3 signal in a group of subepithelial cells in the control animals upon GIPI. Full arrow – caspase-3 signal in the epithelial cells resembling enteroendocrine cells in the STZ-icv animals. Raw data (left), main effects model (upper left) and interaction models (lower left) were presented for all models (A,B,C,D). CAS-3 – caspase-3; GIP – glucose-dependent insulinotropic polypeptide; GIPI – GIP inhibitor [Pro3]-GIP; STZ – streptozotocin; CTR – control; SAL – saline.

### The association of epithelial apoptosis with intestinal morphometric parameters

Finally, the association between activation of epithelial caspase-3 and morphometric parameters was examined utilizing the principal component analysis (**Fig 4**). The first component captured 56.9% of the variation from the data while the second retained 27.2%. Activation of apoptosis reflected by nuclear caspase-3 signal clustered with the villus-related morphometric parameters (VL/CD and VL) (**Fig 4B**).

**Fig 4.**
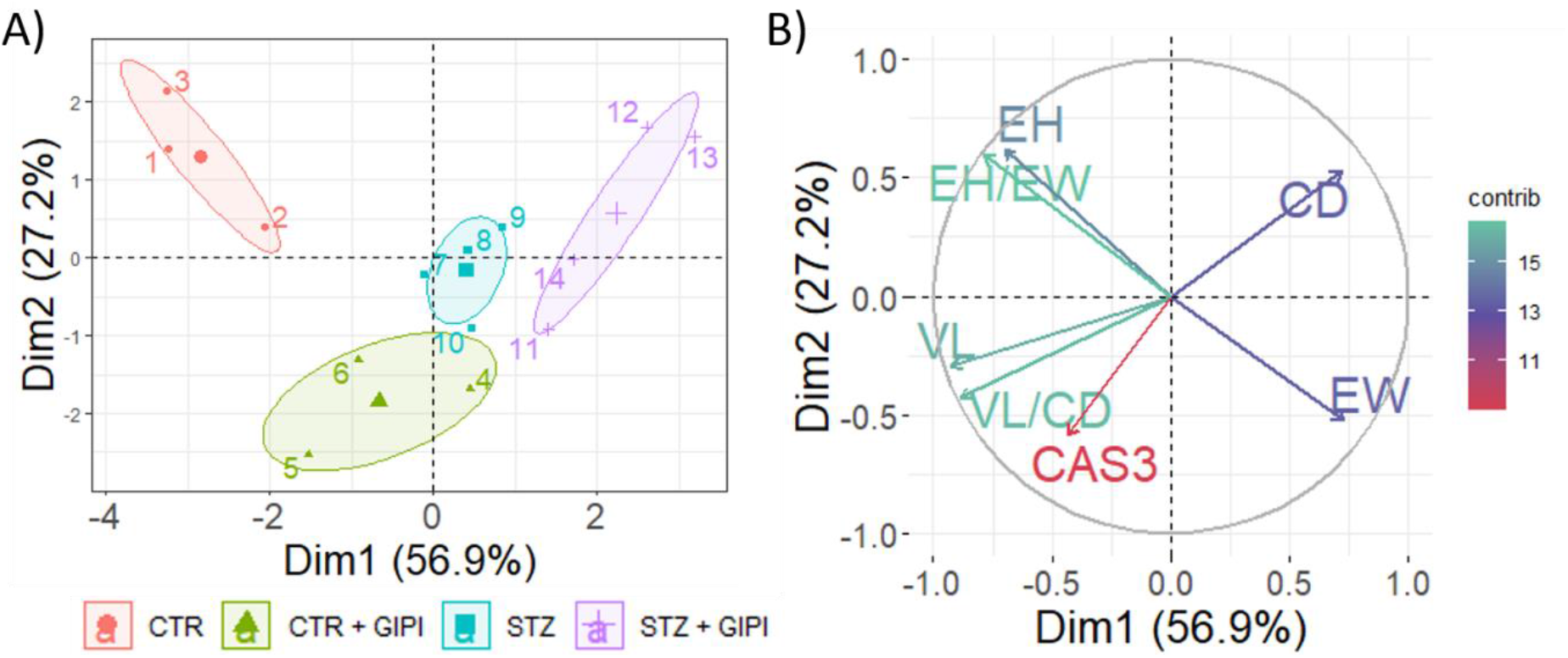
Principal component analysis of epithelial nuclear caspase-3 signal and morphometric parameters in the duodenum of the control rats and the rat model of sporadic Alzheimer’s disease induced by intracerebroventricular streptozotocin (STZ-icv) with and without treatment with intracerebroventricular glucose-dependent insulinotropic polypeptide receptor inhibitor [Pro3]-GIP. **(A)** Individual animals are shown in respect to the biplot. **(H)** Contribution of individual variables in respect to the biplot. Dim 1 –1^st^ dimension; Dim 2 – 2^nd^ dimension; CTR – control animals; STZ – animals treated intracerebroventricularly with streptozotocin; CTR + GIPI – the control animals treated acutely with intracerebroventricular [Pro^3^]-GIP; STZ + GIPI – STZ animals treated acutely with intracerebroventricular [Pro^3^]-GIP. EH – epithelial cell height; EW – epithelial cell width; CD – crypt depth; VL – villus length; VL/CD – the ratio of villus length and adjacent crypt depth; CAS3 – the intensity of nuclear caspase-3 signal corresponding to activated caspase. contrib – contribution.

## Discussion

The presented results **i)** suggest dysfunctional intestinal epithelial cell turnover and apoptosis in the STZ-icv rat model of sAD; **ii)** extend the current knowledge on the brain-gut axis-mediated control of the intestinal redox homeostasis by indicating that, besides brain GLP-1Rs [25], GIP-Rs are also involved in the control of intestinal reactive oxygen species (ROS) generation and clearance and demonstrate that the brain-gut axis dyshomeostasis in the STZ-icv animals does not only involve brain GLP-1Rs[25], but also brain GIP-Rs, and **iii)** suggest that the brain GIP-R might be involved in the regulation of mechanisms responsible for the maintenance of the GI barrier.

### Dysfunctional intestinal epithelial cell turnover and apoptosis in the duodenum of the STZ-icv rats

The maintenance of the functional GI barrier prevents the intestinal luminal microorganisms, inflammatory mediators, and Aβ from entering into the circulation and triggering inflammatory processes driving systemic inflammation, metabolic dysfunction, the permeability of the blood-brain barrier (BBB), and neurodegeneration[57]. Although the failure of the intestinal mucosal barrier has been reported in the transgenic models of AD (e.g. Tg2576 [19]), the integrity of the GI barrier has never been examined in the non-transgenic models of AD such as the STZ-icv model. We have recently reported that markers of oxidative stress are increased in the duodenum of the STZ-icv rats[25]. One month after the STZ-icv treatment lipid peroxidation was increased and endogenous molecules involved in the thiol-disulfide exchange, namely free sulfhydryl groups of proteins and low molecular weight thiols were consumed in the duodenum of the rat model of sAD [25]. Furthermore, the activity of catalase and superoxide dismutase (SOD), major constituents of the cellular enzymatic antioxidant defense system were reduced in the STZ-icv rats [25]. The unchanged nitrocellulose redox permanganometry-assessed [58] residual reductive capacity suggested the consumption of the endogenous antioxidants was able to at least partially compensate the burden of ROS [25].

In this experiment, we measured plasma and duodenal TBARS to confirm or reject our previous observations [25]. Interestingly, while the values of plasma TBARS indicated no significant difference between the STZ-icv animals and the controls in our previous experiment [25], here we observed a slight reduction in the STZ-icv group (**Fig 1A-C**). Furthermore, a reduction of plasma TBARS was observed upon inhibition of the brain GIP-R in both the controls and the rat model of sAD similarly as has been observed following the inhibition of GLP-1R [25], however here the effect was more pronounced (**Fig 1A-C**).

The presence of duodenal oxidative stress in the STZ-icv rats was confirmed by the findings of increased lipid peroxidation in an independent cohort of the STZ-icv rats (**Fig 1D-F**). Oxidative stress has been recognized as an important contributor to the disruption of the epithelial tight junctions and ROS-mediated GI permeability has been shown to play a crucial role in the pathogenesis of GI disorders such as inflammatory bowel disease, alcoholic endotoxemia, infectious enterocolitis, celiac disease, and necrotizing enterocolitis [59]. Recent evidence suggests ROS-induced GI permeability might also be involved in the pathogenesis and progression of AD and dysbiosis has been proposed as a potential source of intestinal oxidative stress [60]. The intestinal function is reflected in its morphology and morphometric analysis of the gut can provide valuable insight into GI homeostasis [61]. VL/CD is an especially interesting and informative parameter as it reflects epithelial cell turnover, an integral mechanism involved in the maintenance of the GI barrier [55]. The rat upper small intestinal VL/CD decreases with age, and it has been proposed that this process reflects the observed age-related impairment of the proximal gut absorptive capacity[62]. Furthermore, the decreased VL/CD accompanies the development of other pathophysiological processes such as dysbiosis and intestinal oxidative stress in different animal models of intestinal dyshomeostasis (e.g. heat-stress-induced intestinal damage in broilers [63], glyphosate [64], or high salt-fed rats [65]). In the STZ-icv-treated rats, VL and VL/CD were found to be decreased, and CD was increased one month after the treatment indicating increased duodenal oxidative stress is accompanied by a disbalance of epithelial cell turnover (**Fig 2A-C**). Considering the contraction of the villus core has been recognized as an important mechanism that minimizes the area of the denuded basement membrane that needs to be re-epithelized[56,66], the shorter villi (**Fig 2B**) and decreased VL/CD (**Fig 2A**) suggest that enterocyte loss from the villus exceeded the regenerative capacity of the crypts in the STZ-icv rats. The contraction of the villus core during the process of reepithelization is accompanied by morphometric adaptation of the remaining epithelial cells that flatten to cover the denuded area of the basement membrane [56]. The remaining epithelial cells covering the villus tip were shorter in the STZ-icv rats, and the distances between adjacent epithelial nuclei were greater suggesting activation of the mechanisms involved in the repair of the intestinal barrier (**Fig 2D-F**).

The intestinal epithelial cell shedding has been recognized as a highly complex mechanism in which cell turnover has to take place without undermining the integrity of the mucosal barrier. Considering the failure of the physiological regulation of the epithelial cell turnover leads to pathological shedding that leaves the GI barrier permeable to luminal microorganisms, noxious substances, and enzymes promoting systemic and neuro-inflammation, the loss of integrity of the GI barrier in the STZ-icv rats could potentiate the development of the AD-like neuropathological changes (**Fig 5**).

**Fig 5.**
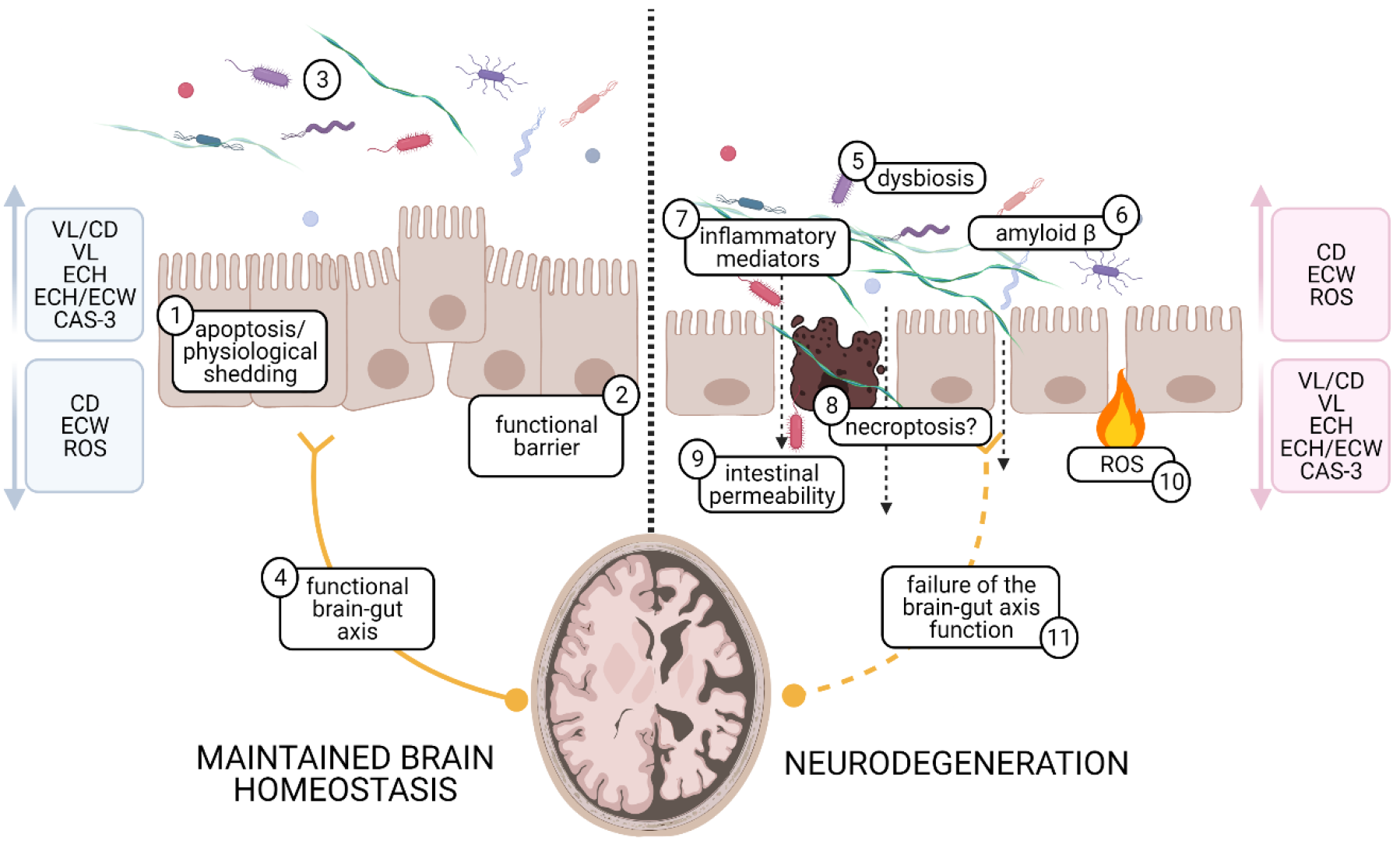
A schematic representation of the potential mechanisms by which failure of the physiological regulation of the epithelial cell turnover might potentiate the development of neuroinflammation and neurodegeneration. **(1)** Apoptosis and physiological shedding maintain the integrity and **(2)** function of the gastrointestinal barrier. **(3)** Healthy gut microbiota promotes the maintenance of gut homeostasis and prevents the development of inflammation. **(4) The** Brain-gut axis maintains gut homeostasis by regulating permeability, motility, secretion, the immune function of the gut, and the host-enteric microbiota signaling. **(5)** Intestinal dysbiosis promotes systemic immune-metabolic dyshomeostasis by secretion of **(6)** amyloid β and **(7)** proinflammatory mediators. Disbalance of the epithelial cell turnover and apoptosis lead to activation of **(8)** alternative epithelial pro-inflammatory cell death pathways (e.g. necroptosis) and development of **(9)** intestinal permeability. **(10)** Redox disbalance promotes the development of intestinal permeability by undermining intestinal homeostasis. Failure of gut homeostasis and intestinal barrier enable the breach of luminal microorganisms, inflammatory mediators, and amyloid β into the systemic circulation and stimulate the development of systemic immune-metabolic dyshomeostasis and subsequently neuroinflammation. **(11)** Failure of the brain-gut axis function promotes the development of intestinal pathophysiology via deficient regulation of permeability, motility, secretion, the immune function of the gut, and the host-enteric microbiota signaling. EH – epithelial cell height; EW – epithelial cell width; CD – crypt depth; VL – villus length; VL/CD – the ratio of villus length and adjacent crypt depth; CAS3 – caspase signaling regulating apoptosis; ROS – reactive oxygen species.

To further explore the disbalance of the intestinal epithelial turnover in the STZ-icv rats, we focused on caspase-3 (**Fig 3**) – a key effector of caspase mediating apoptosis[67]. Both the western blots and the multiplex fluorescent signal amplification indicated reduced caspase-3 in the duodenum of the STZ-icv rats (**Fig 3**). The concentration of both the 17 and 30 kDa caspase-3 fragments were found to be reduced in the duodenal homogenates of the STZ-icv animals (**Fig 3A,B**) indicating that the suppression of apoptosis might be mediated by the reduced expression of the inactive proenzyme rather than inhibition of its proteolytic activation as the active-to-inactive fragment ratio remained largely unchanged by the STZ-icv treatment (**Fig 3C**). A similar trend was observed in the duodenal epithelial cells upon multiplex fluorescent signal amplification (**Fig 3D,E**). Analysis of the association of the caspase-3 activation with the morphometric parameters indicated that apoptosis was associated with increased VL and VL/CD (**Fig 4**).

Several major cell death pathways are involved in the processes regulating intestinal epithelial turnover[68]. Apoptosis and anoikis have been recognized as the most important intestinal epithelial programmed cell death pathways that mediate the removal of dying cells without eliciting inflammatory responses[68]. In contrast, pyroptosis and necroptosis are involved in programmed cell death processes in which inflammatory signaling is activated[68]. Pyroptosis is often triggered by pathogen-induced activation of the inflammasome, and necroptosis is activated when programmed cell death signaling is activated, but the caspase cascade fails due to the inactivation of the effector caspases[68]. Importantly, in the intestinal epithelium, apoptosis and anoikis-mediated epithelial shedding maintain the integrity of the GI barrier while pyroptosis and necroptosis enable enterocyte contents, pathogens, and pro-inflammatory molecules to leak out into the surrounding tissue and trigger inflammatory response[68]. To the best of our knowledge, intestinal apoptosis has never been examined in the STZ-icv rats. The suppression of apoptosis and reduced activation of caspase-3 in the small intestine have been reported in the intravenous streptozotocin-induced rat model of diabetes[69]. Interestingly, in contrast to hypoplasia observed in the STZ-icv rats (**Fig 2**), inhibition of intestinal apoptosis was accompanied by mucosal hyperplasia in streptozotocin-induced diabetes[69]. Taken together, these findings indicate that inhibition of apoptosis led to a decrease in epithelial cell death in streptozotocin-induced diabetes but failed to suppress cell death in the STZ-icv rat model of sAD. The exact underlying mechanisms are to be explored, however, it is possible that alternative pathways of cell death were induced in the STZ-icv model. One possibility is that necroptotic pathways are activated in the duodenum of the STZ-icv rats as necroptosis occurs under conditions in which apoptotic effector caspases are inhibited or inactive and external stimuli activate the tumor necrosis factor receptors and/or toll-like receptors (TLR) to phosphorylate the mixed linked kinase-like protein MLKL[68]. In this scenario, a shift from the apoptotic to the necroptotic epithelial cell death could result in the disruption of the GI-barrier and maintenance of oxidative stress in the intestine of the STZ-icv rats driving systemic and neuro-inflammation. Even if the suppression of epithelial apoptosis was not directly involved in the primary intestinal pathophysiological processes in the STZ-icv rats, the state of inhibited epithelial caspase cascade would make the intestine susceptible to the activation of the vicious self-sustaining inflammatory cycle in the presence of noxious stimuli activating the TLRs [70] (e.g. dysbiosis[71] or circadian dysrhythmia[72]). Whether the inhibition of apoptosis in the STZ-icv potentiates intestinal epithelial necroptosis remains to be explored in our future experiments.

Whether the STZ-icv treatment affects all subpopulations of intestinal epithelial cells equally is another important question to be answered in the upcoming experiments. The disbalance of intestinal goblet cell turnover and function might affect the GI barrier by affecting mucus production[11]. Peripherally administered streptozotocin was found to be toxic towards neuroendocrine cells of the gut[73], however, the effect on the population of intestinal neuroendocrine cells was never examined in the STZ-icv rat model of sAD. Considering the plasma active GLP-1 was found to be decreased in the STZ-icv animals 3 months after the model induction[30], it is possible that the observed effect is a consequence of its action on the turnover or function of gut neuroendocrine cells. Further exploration of specific subpopulations of intestinal cells affected by the STZ-icv and their role in the development of the GI pathophysiology might provide an important insight into the pathogenesis of neurodegenerative processes in the rat model of sAD.

### Inhibition of the brain GIP-R induces duodenal redox dyshomeostasis in the control, but not in the STZ-icv rats

We have recently reported that the inhibition of the brain GLP-1Rs by intracerebroventricular administration of exendin-3(9-39)amide can affect duodenal redox homeostasis and induce oxidative stress in the control rats[25]. Interestingly, the rats treated with the STZ-icv 1 month before were resistant to the brain GLP-1R inhibition-mediated induction of redox dyshomeostasis[25]. The exact nature of the phenomenon remains to be explored. The two most obvious explanations are that: **i)** increased baseline intestinal oxidative stress masks the effect in the STZ-icv rat model of sAD, or **ii)** the brain-gut axis of the STZ-icv animals is resistant to the perturbation of incretin signaling in the brain. Nevertheless, the pro-oxidative effects of inhibition of the brain GLP-1R signaling were evident in the plasma of the STZ-icv rats despite the increased baseline oxidative stress[25]. Furthermore, some of the redox-related effects on the duodenum were characterized by qualitative interactions rather than the absence of the effect indicating that the observed differences are a result of the dysfunctional incretin effect rather than the concealment by baseline oxidative stress levels [25]. Here, we provide additional information on the brain-gut incretin axis by demonstrating that inhibition of brain GIP-R signaling acutely induces duodenal oxidative stress in the control rats (**Fig 1D-F**) in a similar manner as was reported upon inhibition of the brain GLP-1R[25]. Interestingly, the STZ-icv rats were resistant to the effect indicating that the phenomenon observed in previous research [25] is not specific for GLP-1, but that the STZ-icv duodenal redox homeostasis is also resistant to the inhibition of the brain GIP-Rs (**Fig 1D-F**). Considering that a similar effect of brain incretin signaling-inhibition-mediated duodenal oxidative stress was observed in two separate cohorts of control animals and that two separate cohorts of the STZ-icv animals were resistant to the induction of duodenal redox dyshomeostasis by inhibition of different incretin receptors in the brain, we propose that the observed effects might be mediated by a common mechanism.

### Involvement of the brain GIP-R in the regulation of intestinal barrier maintenance

The analysis of duodenal morphometric parameters was primarily done to explore whether the gut oxidative stress was associated with abnormal morphology of the GI barrier in the STZ-icv rats. Nevertheless, some unexpected findings related to the effects of acute pharmacological inhibition of the brain GIP-R were made. Two observations that were particularly surprising were that **i)** the inhibition of brain GIP-R was able to significantly affect duodenal morphological parameters just 30 minutes after the treatment, and **ii)** in the STZ-icv animals, inhibition of the brain GIP-R primarily affected the morphology of the villi, while in the controls the effect on the individual epithelial cells was more pronounced (**Fig 2**). Morphometric parameters do not only reflect chronic structural changes in tissue cytoarchitectonics, but also its functional characteristics. The villus contraction occurs immediately after mucosal injury and it plays an integral role in the process of reepithelization as it minimizes the area of the denuded basement membrane[66,74]. The exact mechanisms of contraction are still being explored, but myofibroblasts have been shown to play a key role and the process seems to be energetically demanding and regulated by the nervous system as dinitrophenol-induced depletion of ATP as well as tetrodotoxin denervation both inhibit acute shortening of the villi upon intestinal injury[66,74]. Taking this into account, the morphometric changes induced upon acute pharmacological inhibition of cerebral GIP-Rs (**Fig 2**) more likely reflect acute epithelial repair mechanisms than tissue cytoarchitectonic structure. This phenomenon remains to be further explored, however rapid onset of the effect suggests the brain GIP-Rs might be involved in the physiological regulation of the intestinal barrier function via the brain-gut axis. The observation that the effect of the intracerebroventricular [Pro^3^]-GIP was more pronounced in the individual epithelial cells in the control animals (**Fig 2D-F**), in contrast to the change in villus structure observed primarily in the STZ-icv rats (**Fig 2A-C**) indicates that the brain GIP-Rs might control intestinal integrity by more than one mechanism. Another potential explanation might be that the absence of further epithelial morphometric adaptation to the disruption of intestinal integrity by inhibition of cerebral GIP-Rs in the STZ-icv rats might simply reflect maximal saturation of the mucosal repair mechanisms at baseline as similar ECH/ECW has been observed in the STZ-icv rats not treated with [Pro^3^]-GIP (**Fig 2**). In line with this, the morphometric adaptation of the villus might be a part of a rescue mechanism activated once epithelial morphometric adaptation cannot fully compensate mucosal dyshomeostasis and maintain barrier integrity. This explanation remains highly hypothetical, however, if this is the case, morphometric adaptation of individual epithelial cells upon intracerebroventricular [Pro^3^]-GIP might be enough to maintain intestinal integrity and prevent the activation of a more robust villus response in the controls (**Fig 2**). At the same time, such staircase phenomenon might explain why intracerebroventricular [Pro^3^]-GIP activates repair mechanisms at the level of the villus in the STZ-icv rats where adaptory response at the level of individual epithelial cells is already fully exploited (**Fig 2**). Further dissection of the role of brain GIP-R might provide a novel insight into mucosal repair mechanisms and elucidate whether GIP-R might be exploited as a potential therapeutic target in the context of mucosal repair.

## Conclusions

The role of the GI system in the etiopathogenesis and the progression of neurodegeneration is being actively explored both in humans and in the animal models of AD. Considering non-transgenic models of AD provide a unique insight into pathomechanisms mediating the effects of risk factors on the progression of the disease understanding the pathophysiological processes in the GI tract of the non-transgenic models might provide an important platform for designing new preventive, diagnostic, and treatment strategies. Here we confirm the presence of oxidative stress in the duodenum of the STZ-icv rat model of sAD and provide further insights into the GI-related pathophysiological processes reflected by morphometric abnormalities and dysfunctional duodenal epithelial apoptosis. The observed disbalance of the duodenal epithelial cell turnover and apoptosis may undermine the integrity of the GI barrier and promote systemic and neuro-inflammatory processes driving neurodegeneration. Furthermore, the duodenal redox homeostasis was found to be similarly insensitive to brain GIP-R inhibition as was reported for the brain GLP-1R suggesting a general resistance of the brain-gut incretin axis might be present and possibly involved in the pathophysiological processes induced by the STZ-icv. Finally, involvement of the brain GIP-R in the regulation of intestinal mucosal repair mechanisms has been reported warranting further investigation of the brain-gut incretin axis for potential pharmacological exploitation.

## Author’s contributions

**JH, ABP, AK,** and **JOB** conducted the *in vivo* part of the experiment. **JH** prepared the samples, performed plasma and duodenal TBARS measurement, western blot, morphometric analyses, designed multiplex fluorescent signal amplification and the apparatus for the elimination of autofluorescence, synthesized fluorescent tyramide conjugates, and conducted CARD. **JH** conducted data curation, analysis, and visualization and wrote the manuscript. **ABP**, **AK**, **JOB**, **FK**, **CS**, R**PR**, and **MSP** commented on the manuscript and provided critical feedback. **MSP**(mentor of JH and PI of the lab) supervised the project and provided funding.

## Data and code availability

A complete dataset and code used for the analysis are available upon request to the corresponding author.

## Ethics committee approval

All experiments were conducted in concordance with the highest standard of animal welfare. Only certified personnel handled animals. Animal procedures were carried out at the University of Zagreb Medical School (Zagreb, Croatia) and complied with current institutional, national (The Animal Protection Act, NN 102/17; NN32/19), and international (Directive 2010/63/EU) guidelines governing the use of experimental animals. The experiments were approved by the national regulatory body responsible for issuing ethical approvals, the Croatian Ministry of Agriculture, and the Ethical Committee of the University of Zagreb School of Medicine.

## References

1. Mayeux R, Stern Y. Epidemiology of Alzheimer Disease. Cold Spring Harb Perspect Med [Internet]. 2012 [cited 2021 Apr 1];2. Available from: https://www.ncbi.nlm.nih.gov/pmc/articles/PMC3405821/

2. Bekris LM, Yu C-E, Bird TD, Tsuang DW. Genetics of Alzheimer Disease. J Geriatr Psychiatry Neurol [Internet]. 2010 [cited 2021 Feb 24];23:213–27. Available from: https://www.ncbi.nlm.nih.gov/pmc/articles/PMC3044597/

3. Fu P, Gao M, Yung KKL. Association of Intestinal Disorders with Parkinson’s Disease and Alzheimer’s Disease: A Systematic Review and Meta-Analysis. ACS Chem Neurosci. 2020;11:395–405.

4. Jiang C, Li G, Huang P, Liu Z, Zhao B. The Gut Microbiota and Alzheimer’s Disease. J Alzheimers Dis. 2017;58:1–15.

5. Angelucci F, Cechova K, Amlerova J, Hort J. Antibiotics, gut microbiota, and Alzheimer’s disease. J Neuroinflammation. 2019;16:108.

6. Carabotti M, Scirocco A, Maselli MA, Severi C. The gut-brain axis: interactions between enteric microbiota, central and enteric nervous systems. Ann Gastroenterol. 2015;28:203–9.

7. Rhee SH, Pothoulakis C, Mayer EA. Principles and clinical implications of the brain–gut–enteric microbiota axis. Nat Rev Gastroenterol Hepatol [Internet]. 2009 [cited 2021 Apr 6];6. Available from: https://www.ncbi.nlm.nih.gov/pmc/articles/PMC3817714/

8. Homolak J, Mudrovčić M, Vukić B, Toljan K. Circadian Rhythm and Alzheimer’s Disease. Med Sci (Basel) [Internet]. 2018 [cited 2021 Apr 1];6. Available from: https://www.ncbi.nlm.nih.gov/pmc/articles/PMC6164904/

9. Teichman EM, O’Riordan KJ, Gahan CGM, Dinan TG, Cryan JF. When Rhythms Meet the Blues: Circadian Interactions with the Microbiota-Gut-Brain Axis. Cell Metabolism [Internet]. 2020 [cited 2021 Apr 1];31:448–71. Available from: https://www.sciencedirect.com/science/article/pii/S155041312030067X

10. Segers A, Depoortere I. Circadian clocks in the digestive system. Nature Reviews Gastroenterology & Hepatology [Internet]. Nature Publishing Group; 2021 [cited 2021 Apr 1];18:239–51. Available from: https://www.nature.com/articles/s41575-020-00401-5

11. Herath M, Hosie S, Bornstein JC, Franks AE, Hill-Yardin EL. The Role of the Gastrointestinal Mucus System in Intestinal Homeostasis: Implications for Neurological Disorders. Front Cell Infect Microbiol [Internet]. Frontiers; 2020 [cited 2021 Apr 1];10. Available from: https://www.frontiersin.org/articles/10.3389/fcimb.2020.00248/full

12. Simic G, Stanic G, Mladinov M, Jovanov-Milosevic N, Kostovic I, Hof PR. Does Alzheimer’s disease begin in the brainstem? Neuropathol Appl Neurobiol. 2009;35:532–54.

13. Polak T, Ehlis A-C, Langer JBM, Plichta MM, Metzger F, Ringel TM, et al. Non-invasive measurement of vagus activity in the brainstem – a methodological progress towards earlier diagnosis of dementias? J Neural Transm (Vienna). 2007;114:613–9.

14. Leblhuber F, Ehrlich D, Steiner K, Geisler S, Fuchs D, Lanser L, et al. The Immunopathogenesis of Alzheimer’s Disease Is Related to the Composition of Gut Microbiota. Nutrients [Internet]. 2021 [cited 2021 Apr 1];13. Available from: https://www.ncbi.nlm.nih.gov/pmc/articles/PMC7912578/

15. Soto M, Herzog C, Pacheco JA, Fujisaka S, Bullock K, Clish CB, et al. Gut microbiota modulate neurobehavior through changes in brain insulin sensitivity and metabolism. Molecular Psychiatry [Internet]. Nature Publishing Group; 2018 [cited 2021 Apr 1];23:2287–301. Available from: https://www.nature.com/articles/s41380-018-0086-5

16. Friedland RP, Chapman MR. The role of microbial amyloid in neurodegeneration. PLoS Pathog. 2017;13:e1006654.

17. Brown GC. The endotoxin hypothesis of neurodegeneration. J Neuroinflammation. 2019;16:180.

18. Sun Y, Sommerville NR, Liu JYH, Ngan MP, Poon D, Ponomarev ED, et al. Intra-gastrointestinal amyloid-β1-42 oligomers perturb enteric function and induce Alzheimer’s disease pathology. J Physiol. 2020;598:4209–23.

19. Honarpisheh P, Reynolds CR, Blasco Conesa MP, Moruno Manchon JF, Putluri N, Bhattacharjee MB, et al. Dysregulated Gut Homeostasis Observed Prior to the Accumulation of the Brain Amyloid-β in Tg2576 Mice. Int J Mol Sci. 2020;21.

20. Semar S, Klotz M, Letiembre M, Van Ginneken C, Braun A, Jost V, et al. Changes of the enteric nervous system in amyloid-β protein precursor transgenic mice correlate with disease progression. J Alzheimers Dis. 2013;36:7–20.

21. Brandscheid C, Schuck F, Reinhardt S, Schäfer K-H, Pietrzik CU, Grimm M, et al. Altered Gut Microbiome Composition and Tryptic Activity of the 5xFAD Alzheimer’s Mouse Model. J Alzheimers Dis. 2017;56:775–88.

22. Chi H, Cao W, Zhang M, Su D, Yang H, Li Z, et al. Environmental noise stress disturbs commensal microbiota homeostasis and induces oxi-inflammmation and AD-like neuropathology through epithelial barrier disruption in the EOAD mouse model. J Neuroinflammation. 2021;18:9.

23. Wang Y, An Y, Ma W, Yu H, Lu Y, Zhang X, et al. 27-Hydroxycholesterol contributes to cognitive deficits in APP/PS1 transgenic mice through microbiota dysbiosis and intestinal barrier dysfunction. J Neuroinflammation. 2020;17:199.

24. Sohrabi M, Pecoraro HL, Combs CK. Gut Inflammation Induced by Dextran Sulfate Sodium Exacerbates Amyloid-β Plaque Deposition in the AppNL-G-F Mouse Model of Alzheimer’s Disease. J Alzheimers Dis. 2021;79:1235–55.

25. Homolak J, Babic Perhoc A, Knezovic A, Osmanovic Barilar J, Salkovic-Petrisic M. Failure of the brain glucagon-like peptide-1-mediated control of intestinal redox homeostasis in a rat model of sporadic Alzheimer′s disease. bioRxiv [Internet]. 2021 [cited 2021 Mar 25]; Available from: https://www.biorxiv.org/content/10.1101/2021.03.22.436453v1

26. Akbarzadeh A, Norouzian D, Mehrabi MR, Jamshidi Sh, Farhangi A, Verdi AA, et al. Induction of diabetes by Streptozotocin in rats. Indian J Clin Biochem [Internet]. 2007 [cited 2021 Apr 2];22:60–4. Available from: https://www.ncbi.nlm.nih.gov/pmc/articles/PMC3453807/

27. Chao P-C, Li Y, Chang C-H, Shieh JP, Cheng J-T, Cheng K-C. Investigation of insulin resistance in the popularly used four rat models of type-2 diabetes. Biomedicine & Pharmacotherapy [Internet]. 2018 [cited 2021 Apr 2];101:155–61. Available from: https://www.sciencedirect.com/science/article/pii/S0753332217363801

28. Correia SC, Santos RX, Perry G, Zhu X, Moreira PI, Smith MA. Insulin-resistant brain state: the culprit in sporadic Alzheimer’s disease? Ageing Res Rev. 2011;10:264–73.

29. Barilar JO, Knezovic A, Perhoc AB, Homolak J, Riederer P, Salkovic-Petrisic M. Shared cerebral metabolic pathology in non-transgenic animal models of Alzheimer’s and Parkinson’s disease. J Neural Transm [Internet]. 2020 [cited 2021 Apr 2];127:231–50. Available from: https://doi.org/10.1007/s00702-020-02152-8

30. Knezovic A, Osmanovic Barilar J, Babic A, Bagaric R, Farkas V, Riederer P, et al. Glucagon-like peptide-1 mediates effects of oral galactose in streptozotocin-induced rat model of sporadic Alzheimer’s disease. Neuropharmacology. 2018;135:48–62.

31. Sharma M, Gupta YK. Intracerebroventricular injection of streptozotocin in rats produces both oxidative stress in the brain and cognitive impairment. Life Sciences [Internet]. 2001 [cited 2021 Feb 25];68:1021–9. Available from: https://www.sciencedirect.com/science/article/pii/S0024320500010055

32. Ghosh R, Sil S, Gupta P, Ghosh T. Optimization of intracerebroventricular streptozotocin dose for the induction of neuroinflammation and memory impairments in rats. Metab Brain Dis [Internet]. 2020 [cited 2021 Feb 25];35:1279–86. Available from: https://doi.org/10.1007/s11011-020-00588-1

33. Knezovic A, Loncar A, Homolak J, Smailovic U, Osmanovic Barilar J, Ganoci L, et al. Rat brain glucose transporter-2, insulin receptor and glial expression are acute targets of intracerebroventricular streptozotocin: risk factors for sporadic Alzheimer’s disease? J Neural Transm (Vienna). 2017;124:695–708.

34. Blokland A, Jolles J. Spatial learning deficit and reduced hippocampal ChAT activity in rats after an ICV injection of streptozotocin. Pharmacol Biochem Behav. 1993;44:491–4.

35. Salkovic-Petrisic M, Osmanovic-Barilar J, Brückner MK, Hoyer S, Arendt T, Riederer P. Cerebral amyloid angiopathy in streptozotocin rat model of sporadic Alzheimer’s disease: a long-term follow up study. J Neural Transm (Vienna). 2011;118:765–72.

36. Li Y, Xu P, Shan J, Sun W, Ji X, Chi T, et al. Interaction between hyperphosphorylated tau and pyroptosis in forskolin and streptozotocin induced AD models. Biomedicine & Pharmacotherapy [Internet]. 2020 [cited 2021 Feb 25];121:109618. Available from: https://www.sciencedirect.com/science/article/pii/S0753332219352400

37. Knezovic A, Osmanovic-Barilar J, Curlin M, Hof PR, Simic G, Riederer P, et al. Staging of cognitive deficits and neuropathological and ultrastructural changes in streptozotocin-induced rat model of Alzheimer’s disease. J Neural Transm [Internet]. 2015 [cited 2021 Feb 26];122:577–92. Available from: https://link.springer.com/article/10.1007/s00702-015-1394-4

38. Hölscher C. Insulin Signaling Impairment in the Brain as a Risk Factor in Alzheimer’s Disease. Front Aging Neurosci [Internet]. Frontiers; 2019 [cited 2021 Mar 8];11. Available from: https://www.frontiersin.org/articles/10.3389/fnagi.2019.00088/full

39. Cani PD, Knauf C, Iglesias MA, Drucker DJ, Delzenne NM, Burcelin R. Improvement of Glucose Tolerance and Hepatic Insulin Sensitivity by Oligofructose Requires a Functional Glucagon-Like Peptide 1 Receptor. Diabetes [Internet]. American Diabetes Association; 2006 [cited 2021 Mar 8];55:1484–90. Available from: https://diabetes.diabetesjournals.org/content/55/5/1484

40. MacDonald PE, El-kholy W, Riedel MJ, Salapatek AMF, Light PE, Wheeler MB. The Multiple Actions of GLP-1 on the Process of Glucose-Stimulated Insulin Secretion. Diabetes [Internet]. American Diabetes Association; 2002 [cited 2021 Mar 8];51:S434–42. Available from: https://diabetes.diabetesjournals.org/content/51/suppl_3/S434

41. Li Y, Duffy KB, Ottinger MA, Ray B, Bailey JA, Holloway HW, et al. GLP-1 receptor stimulation reduces amyloid-beta peptide accumulation and cytotoxicity in cellular and animal models of Alzheimer’s disease. J Alzheimers Dis. 2010;19:1205–19.

42. Salcedo I, Tweedie D, Li Y, Greig NH. Neuroprotective and neurotrophic actions of glucagon-like peptide-1: an emerging opportunity to treat neurodegenerative and cerebrovascular disorders. Br J Pharmacol [Internet]. 2012 [cited 2021 Mar 8];166:1586–99. Available from: https://www.ncbi.nlm.nih.gov/pmc/articles/PMC3419902/

43. Perry TA, Greig NH. A new Alzheimer’s disease interventive strategy: GLP-1. Curr Drug Targets. 2004;5:565–71.

44. Hölscher C. Brain insulin resistance: role in neurodegenerative disease and potential for targeting. Expert Opinion on Investigational Drugs [Internet]. Taylor & Francis; 2020 [cited 2021 Apr 2];29:333–48. Available from: https://doi.org/10.1080/13543784.2020.1738383

45. Hölscher C. Novel dual GLP-1/GIP receptor agonists show neuroprotective effects in Alzheimer’s and Parkinson’s disease models. Neuropharmacology. 2018;136:251–9.

46. Noble EP, Wurtman RJ, Axelrod J. A simple and rapid method for injecting H3-norepinephrine into the lateral ventricle of the rat brain. Life Sci. 1967;6:281–91.

47. Homolak J, Perhoc AB, Knezovic A, Osmanovic Barilar J, Salkovic-Petrisic M. Additional methodological considerations regarding optimization of the dose of intracerebroventricular streptozotocin A response to: “Optimization of intracerebroventricular streptozotocin dose for the induction of neuroinflammation and memory impairments in rats” by Ghosh et al., Metab Brain Dis 2020 July 21. Metab Brain Dis. 2021;36:97–102.

48. Homolak J, Babic Perhoc A, Knezovic A, Kodvanj I, Virag D, Osmanovic Barilar J, et al. Is galactose a hormetic sugar? Evidence from rat hippocampal redox regulatory network. bioRxiv. 2021;

49. Prabhakar PV, Reddy UA, Singh SP, Balasubramanyam A, Rahman MF, Indu Kumari S, et al. Oxidative stress induced by aluminum oxide nanomaterials after acute oral treatment in Wistar rats. J Appl Toxicol. 2012;32:436–45.

50. Han S, Cui Y, Helbing DL. Inactivation of Horseradish Peroxidase by Acid for Sequential Chemiluminescent Western Blot. Biotechnology Journal [Internet]. 2020 [cited 2021 Mar 25];15:1900397. Available from: https://onlinelibrary.wiley.com/doi/abs/10.1002/biot.201900397

51. Sun Y, Ip P, Chakrabartty A. Simple Elimination of Background Fluorescence in Formalin-Fixed Human Brain Tissue for Immunofluorescence Microscopy. J Vis Exp. 2017;

52. Duong H, Han M. A multispectral LED array for the reduction of background autofluorescence in brain tissue. Journal of Neuroscience Methods [Internet]. 2013 [cited 2021 Mar 25];220:46–54. Available from: https://www.sciencedirect.com/science/article/pii/S0165027013002896

53. Hopman AH, Ramaekers FC, Speel EJ. Rapid synthesis of biotin-, digoxigenin-, trinitrophenyl-, and fluorochrome-labeled tyramides and their application for In situ hybridization using CARD amplification. J Histochem Cytochem. 1998;46:771–7.

54. Arganda-Carreras I, Kaynig V, Rueden C, Eliceiri KW, Schindelin J, Cardona A, et al. Trainable Weka Segmentation: a machine learning tool for microscopy pixel classification. Bioinformatics. 2017;33:2424–6.

55. Moniello G, Ariano A, Panettieri V, Tulli F, Olivotto I, Messina M, et al. Intestinal Morphometry, Enzymatic and Microbial Activity in Laying Hens Fed Different Levels of a Hermetia illucens Larvae Meal and Toxic Elements Content of the Insect Meal and Diets. Animals [Internet]. Multidisciplinary Digital Publishing Institute; 2019 [cited 2021 Mar 24];9:86. Available from: https://www.mdpi.com/2076-2615/9/3/86

56. Williams JM, Duckworth CA, Burkitt MD, Watson AJM, Campbell BJ, Pritchard DM. Epithelial Cell Shedding and Barrier Function. Vet Pathol [Internet]. 2015 [cited 2021 Apr 4];52:445–55. Available from: https://www.ncbi.nlm.nih.gov/pmc/articles/PMC4441880/

57. Liu S, Gao J, Zhu M, Liu K, Zhang H-L. Gut Microbiota and Dysbiosis in Alzheimer’s Disease: Implications for Pathogenesis and Treatment. Mol Neurobiol [Internet]. 2020 [cited 2021 Apr 2];57:5026–43. Available from: https://doi.org/10.1007/s12035-020-02073-3

58. Homolak J, Kodvanj I, Babic Perhoc A, Virag D, Knezovic A, Osmanovic Barilar J, et al. Nitrocellulose redox permanganometry: a simple method for reductive capacity assessment | bioRxiv [Internet]. 2020 [cited 2020 Dec 22]. Available from: https://www.biorxiv.org/content/10.1101/2020.06.16.154682v1

59. Rao R. Oxidative Stress-Induced Disruption of Epithelial and Endothelial Tight Junctions. Front Biosci [Internet]. 2008 [cited 2021 Apr 2];13:7210–26. Available from: https://www.ncbi.nlm.nih.gov/pmc/articles/PMC6261932/

60. Luca M, Di Mauro M, Di Mauro M, Luca A. Gut Microbiota in Alzheimer’s Disease, Depression, and Type 2 Diabetes Mellitus: The Role of Oxidative Stress. Oxid Med Cell Longev [Internet]. 2019 [cited 2021 Apr 2];2019. Available from: https://www.ncbi.nlm.nih.gov/pmc/articles/PMC6501164/

61. Khan J, Islam MN. Morphology of the Intestinal Barrier in Different Physiological and Pathological Conditions. Histopathology – Reviews and Recent Advances [Internet]. IntechOpen; 2012 [cited 2021 Apr 2]; Available from: https://www.intechopen.com/books/histopathology-reviews-and-recent-advances/morphology-of-the-intestinal-barrier-in-different-physiological-and-pathological-conditions

62. Holt PR, Pascal RR, Kotler DP. Effect of aging upon small intestinal structure in the Fischer rat. J Gerontol. 1984;39:642–7.

63. Santos RR, Awati A, Hil PJR den, Tersteeg-Zijderveld MHG, Koolmees PA, Fink-Gremmels J. Quantitative histo-morphometric analysis of heat-stress-related damage in the small intestines of broiler chickens. Avian Pathology [Internet]. Taylor & Francis; 2015 [cited 2021 Apr 2];44:19–22. Available from: https://doi.org/10.1080/03079457.2014.988122

64. Tang Q, Tang J, Ren X, Li C. Glyphosate exposure induces inflammatory responses in the small intestine and alters gut microbial composition in rats. Environ Pollut. 2020;261:114129.

65. Wang H, Li S, Fang S, Yang X, Feng J. Betaine Improves Intestinal Functions by Enhancing Digestive Enzymes, Ameliorating Intestinal Morphology, and Enriching Intestinal Microbiota in High-salt stressed Rats. Nutrients. 2018;10.

66. Moore R, Carlson S, Madara JL. Villus contraction aids repair of intestinal epithelium after injury. Am J Physiol. 1989;257:G274–283.

67. Porter AG, Jänicke RU. Emerging roles of caspase-3 in apoptosis. Cell Death Differ. 1999;6:99–104.

68. Patankar JV, Becker C. Cell death in the gut epithelium and implications for chronic inflammation. Nature Reviews Gastroenterology & Hepatology [Internet]. Nature Publishing Group; 2020 [cited 2021 Apr 4];17:543–56. Available from: https://www.nature.com/articles/s41575-020-0326-4

69. Noda T, Iwakiri R, Fujimoto K, Yoshida T, Utsumi H, Sakata H, et al. Suppression of apoptosis is responsible for increased thickness of intestinal mucosa in streptozotocin-induced diabetic rats. Metabolism. 2001;50:259–64.

70. Burgueño JF, Abreu MT. Epithelial Toll-like receptors and their role in gut homeostasis and disease. Nature Reviews Gastroenterology & Hepatology [Internet]. Nature Publishing Group; 2020 [cited 2021 Apr 6];17:263–78. Available from: https://www.nature.com/articles/s41575-019-0261-4

71. de Kivit S, Tobin MC, Forsyth CB, Keshavarzian A, Landay AL. Regulation of Intestinal Immune Responses through TLR Activation: Implications for Pro- and Prebiotics. Front Immunol [Internet]. 2014 [cited 2021 Apr 6];5. Available from: https://www.ncbi.nlm.nih.gov/pmc/articles/PMC3927311/

72. Mukherji A, Kobiita A, Ye T, Chambon P. Homeostasis in intestinal epithelium is orchestrated by the circadian clock and microbiota cues transduced by TLRs. Cell. 2013;153:812–27.

73. Brenna O, Qvigstad G, Brenna E, Waldum HL. Cytotoxicity of streptozotocin on neuroendocrine cells of the pancreas and the gut. Dig Dis Sci. 2003;48:906–10.

74. Blikslager AT, Moeser AJ, Gookin JL, Jones SL, Odle J. Restoration of barrier function in injured intestinal mucosa. Physiol Rev. 2007;87:545–64.

