## Supplement 1 for "Disbalance of the intestinal epithelial cell turnover and apoptosis in a rat model of sporadic Alzheimer’s disease"

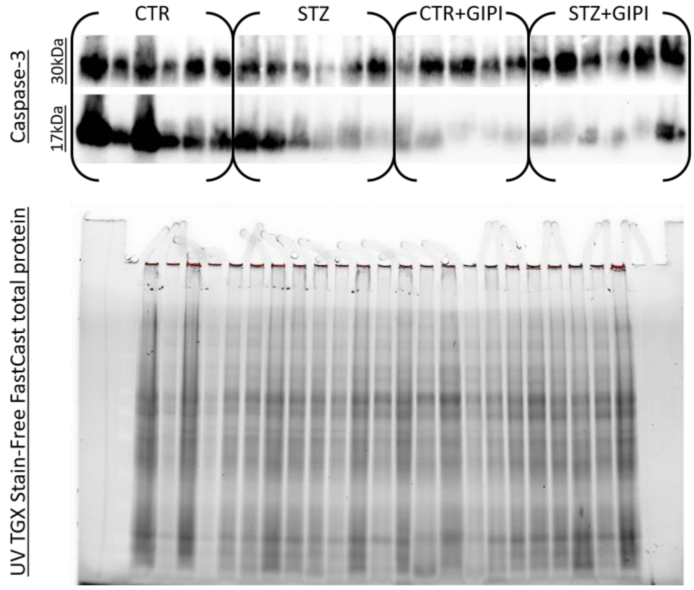


**Fig S1-1. Caspase-3 PAGE and Western blot raw data.**

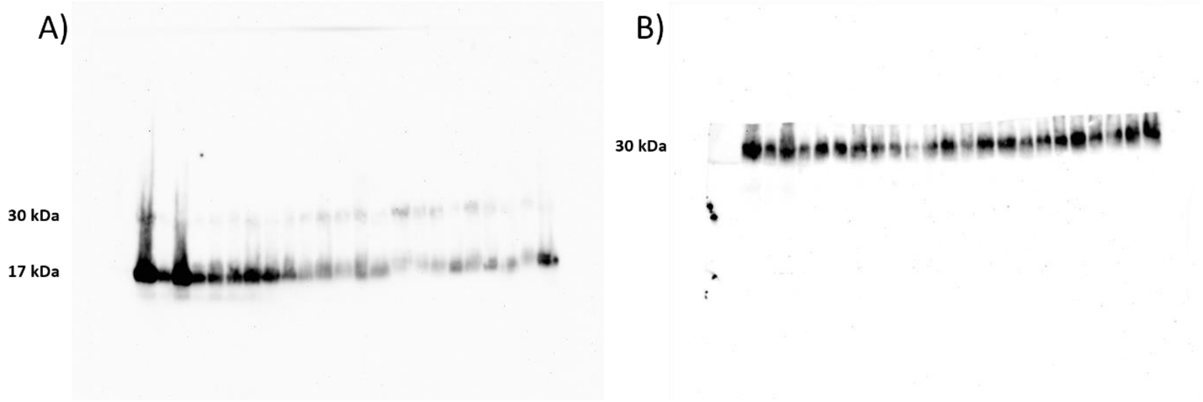


**Fig S1-2. Uncropped images of 17 and 30 kDa caspase-3. (A)** A strong 17 kDa and weak caspaese-3 signal on the same mambrane. **(B)** Lower parto f the membrane was covered by a non-transparent PVC membrane. Upper part of the membrane was cut away.
