## Supplement 2 for "Disbalance of the intestinal epithelial cell turnover and apoptosis in a rat model of sporadic Alzheimer’s disease"

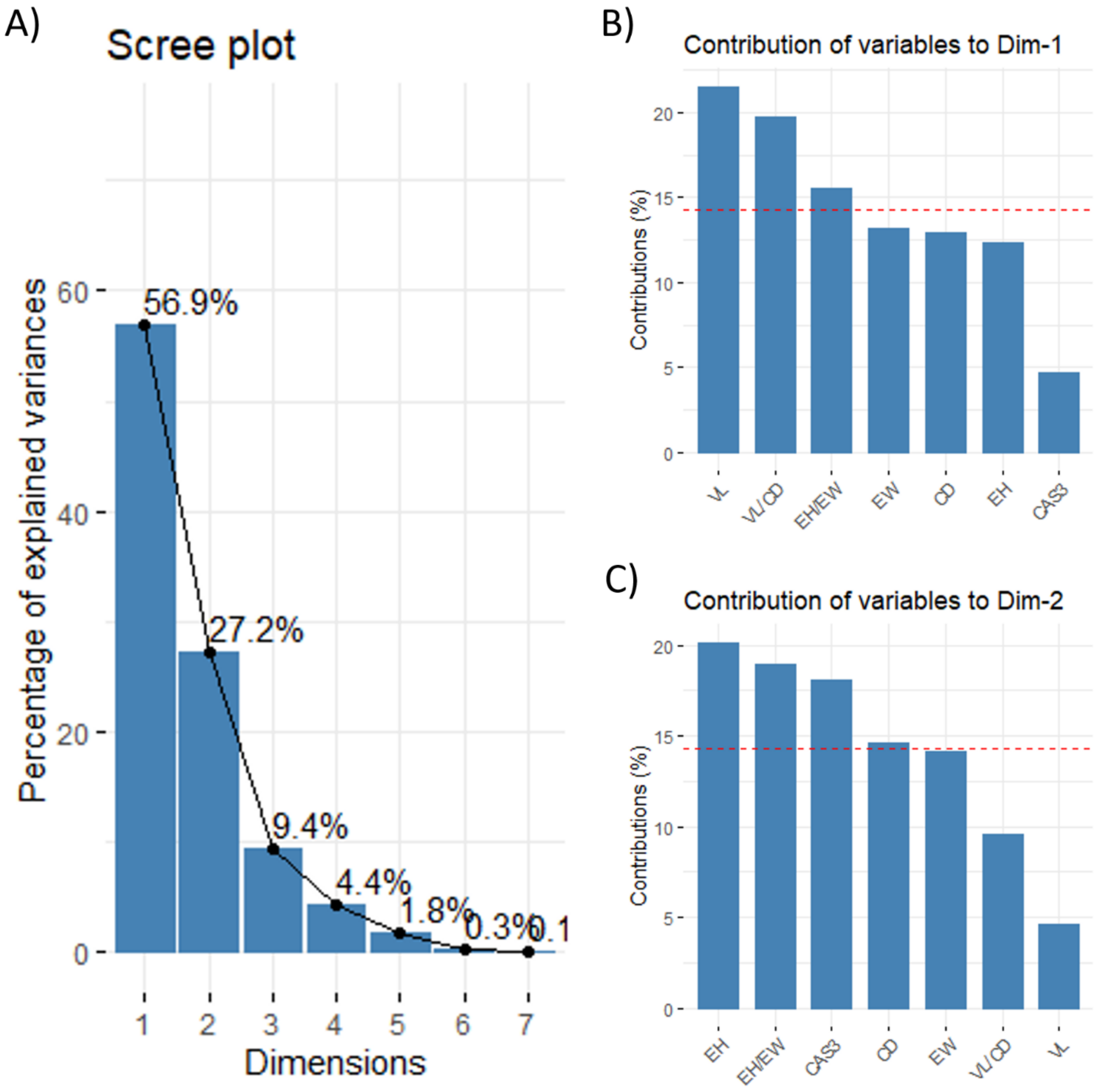


**Fig S2-1.** Principal component analysis **(A)** scree plot, and the contribution of individual variables to the **(B)** first and **(C)** second principal component. Dim 1 –1$^{st}$ dimension; Dim 2 – 2$^{nd}$ dimension; CTR – control animals; STZ – animals treated intracerebroventricularly with streptozotocin; CTR + GIPI – the control animals treated acutely with intracerebroventricular [Pro$^{3}$]-GIP; STZ + GIPI – STZ animals treated acutely with intracerebroventricular [Pro$^{3}$]-GIP. EH – epithelial cell height; EW – epithelial cell width; CD – crypt depth; VL – villus length; VL/CD – the ratio of villus length and adjacent crypt depth; CAS3 – intesnity of nuclear caspase-3 signal corresponding to activated caspase.
